# State anxiety alters the neural oscillatory correlates of predictions and prediction errors during reward learning

**DOI:** 10.1101/2021.03.08.434415

**Authors:** Thomas P Hein, Maria Herrojo Ruiz

## Abstract

Anxiety influences how the brain estimates and responds to uncertainty. These behavioural effects have been described within predictive coding and Bayesian inference frameworks, yet the associated neural correlates remain unclear. Recent work suggests that predictions in generative models of perception are represented in alpha (8–12 Hz) and beta oscillations (13–30 Hz). Updates to predictions are driven by prediction errors weighted by precision (inverse variance; pwPE), and are encoded in gamma oscillations (>30 Hz) and associated with suppression of beta activity. We tested whether state anxiety alters the neural oscillatory activity associated with predictions and pwPE during learning. Healthy human participants performed a probabilistic reward-learning task in a volatile environment. In our previous work, we described learning behaviour in this task using a hierarchical Bayesian model, revealing more precise (biased) beliefs about the reward tendency in state anxiety, consistent with reduced learning in this group. The model provided trajectories of predictions and pwPEs for the current study, allowing us to assess their parametric effects on the time-frequency representations of EEG data. Using convolution modelling for oscillatory responses, we found that, relative to a control group, state anxiety increased beta activity in frontal and sensorimotor regions during processing of pwPE, and in fronto-parietal regions during encoding of predictions. No effects of state anxiety on gamma modulation were found. Our findings expand prior evidence on the oscillatory representations of predictions and pwPEs into the reward-learning domain. The results suggest that state anxiety modulates beta-band oscillatory correlates of pwPE and predictions in generative models, providing insights into the neural processes associated with biased belief updating and poorer reward learning.

## 1. Introduction

Affective states closely interact with decision making (Lerner et al., 2015). For example, altered computations—such as learning rates and estimates of belief uncertainty—during decision making are considered central to explaining clinical conditions including anxiety, depression and stress from a Bayesian predictive coding (Bayesian PC) perspective (Browning et al., 2015; de Berker et al., 2016; Pulcu and Browning, 2019; Williams, 2016). The Bayesian PC framework proposes that the brain continuously updates a hierarchical generative model using predictions optimised through their discrepancy with sensory data—prediction errors (PE)—and weighted by precision (inverse variance; Friston, 2010; Rao and Ballard, 1999; Srinivasan et al., 1982). This hierarchical message passing was hypothesised to be mediated by neural oscillations at specific frequencies, in distinct cortical layers and regions (Bastos et al., 2012). Empirical evidence supports this, identifying predictions in alpha and beta frequencies and PEs in gamma frequencies (Arnal and Giraud, 2012; Auksztulewicz et al., 2017; Bastos et al., 2020; Sedley et al., 2016). Yet how affective states modulate the oscillatory activity associated with predictions and PE signals has been largely overlooked.

Uncertainty makes refining predictions particularly challenging. Estimates of uncertainty (or its inverse, precision) regulate how influential PEs are on updating our generative model of the environment (Friston, 2008; Yu and Dayan, 2005), scaling precision-weighted PEs (pwPEs). Uncertain and changing environments may render prior beliefs obsolete, down-weighting predictions in favour of increasing learning about sensory input. Recent studies have highlighted that precision estimates are important in explaining atypical learning and perception in neuropsychiatric conditions (Fletcher and Frith, 2009; Friston et al., 2013; Lawson et al., 2014; Montague et al., 2012). Anxiety, in particular, has been shown to lead to insufficient adaptation in the face of environmental change (Browning et al., 2015; Huang et al., 2017), disruption in learning and maladaptive biases—in both aversive and reward-learning contexts (Hein et al., 2021; Huang et al., 2017; Kim et al., 2020; Lamba et al., 2020; Piray et al., 2019; Pulcu and Browning, 2019). Whether the learning alterations in anxiety are mediated by oscillatory changes representing predictions and pwPEs remains unknown.

Within the Bayesian PC framework, growing evidence supports that feedforward PE signals are encoded by gamma oscillations (>30 Hz), while backward connections convey predictions expressed in alpha (8–12 Hz) and beta (13–30 Hz) oscillations (Arnal and Giraud, 2012; Bastos et al., 2015; van Pelt et al., 2016; Wang, 2010). Precision weights are also modulated by alpha and beta oscillations (Palmer et al., 2019; Sedley et al., 2016). Because precision weights scale PEs during Bayesian inference and learning (Feldman and Friston, 2010; Friston, 2010), the modulation of pwPE signals could be expressed both in gamma and alpha/beta activity, as recent work suggests (Auksztulewicz et al., 2017). Specifically, gamma increases during pwPE encoding are accompanied by attenuated alpha and beta activity (Auksztulewicz et al., 2017). Indeed, gamma and alpha/beta oscillatory rhythms are anticorrelated across the cortex, as shown in investigations of prediction violations (Bastos et al., 2018; 2020) and during working memory (Lundqvist et al., 2016; 2020; Miller et al., 2018). Complementing these findings, we recently showed that abnormal increases in beta power and burst rate can account for the dampening of pwPEs during reward–based motor learning in anxiety (Sporn et al., 2020).

Accordingly, we speculated that decreased anxiety-related learning during decision making could be associated with abnormally enhanced beta, in addition to reduced gamma, during processing pwPEs.

Predictions have been consistently associated with the modulation of alpha-beta rhythms across multiple modalities, such as visual (Gould et al., 2011), motor (Schoffelen et al., 2005), somatosensory (van Ede et al., 2011), and auditory (Todorovic et al., 2015)—yet frequency-domain evidence for reward-related predictions is currently lacking. Crucially, predictions in deep layers are thought to functionally inhibit the processing of sensory input and PEs in superficial layers (Bastos et al., 2015; Bauer et al., 2014; Mayer et al., 2016; Van Kerkoerle et al., 2014). This suggests that aberrant oscillatory states modulating predictions would be an additional route through which encoding of pwPEs is altered, contributing to impaired learning.

Here, we used convolution modelling of oscillatory responses (Litvak et al., 2013) in previously-acquired EEG data to estimate the neural oscillatory representations of predictions and pwPEs during reward learning in healthy controls and a state anxious group. Our previous computational modelling study (Hein et al., 2021) revealed that state anxiety biases uncertainty estimates, increasing the precision of posterior beliefs about the reward tendency. We now ask whether this bias is associated with altered spectral characteristics of hierarchical message passing, which could represent a candidate marker of biased belief updating and poorer reward learning in anxiety. We hypothesised that the more precise reward predictions found in state anxiety should be associated with increased alpha and beta activity. This, in turn, would inhibit the processing of expected inputs in line with PC accounts, resulting in a hypothesised lower gamma activity and concomitantly higher alpha-beta activity for attenuating encoding of pwPEs.

## 2. Materials & Methods

### 2.1 Participant sample

The data used in the preparation of this work were obtained from our previous study Hein et al. (2021), which was approved by the ethical review committee at Goldsmiths, University of London. Participants were pseudo-randomly allocated into an experimental state anxiety (StA) and control (Cont) group, following a screening phase in which we measured trait anxiety levels in each participant using Spielberger’s Trait Anxiety Inventory (STAI; Spielberger [1983]). Trait anxiety levels were matched in StA and Cont groups: average score and standard error of the mean, SEM: 47 [2.1] in StA, 46 [2.2] in Cont). Importantly, the individual trait anxiety scores were lower than the previously reported clinical level for the general adult population (> 70, Spielberger et al., 1983). Further, the age of the control group (mean 27.7, SEM = 1.2) and their sex (13 female, 8 male) were consistent with those from the state anxiety group (mean 27.5, SEM = 1.3, sex 14 female, 7 male). This is important to consider as there are known age and sex-related confounds to measures of state anxiety (see Voss et al., 2015).

### 2.2 Experimental design

Both groups (StA, Cont) performed a probabilistic binary reward-based learning task where the probability of reward between two images changes across time (Behrens et al., 2007; de Berker et al., 2016; Iglesias et al., 2013). The experiment was divided into four blocks: an initial resting state block (R1: baseline), two reward-learning task blocks (TB1, TB2), and a final resting state block (R2). Each resting state block was 5 minutes. Participants were instructed to relax and keep their eyes open and fixated on a cross in the middle of the presentation screen while we recorded EEG responses from the scalp and EKG responses from the heart.

The experimental task consisted of 200 trials in each task block (TB1, TB2). The aim was for participants to maximise reward across all trials by predicting which of the two images (blue, orange) would reward them (win, positive reinforcement, 5 pence reward) or not (lose, 0 pence reward). The probability governing reward for each stimulus (reciprocal: p, 1−p) changed across the experiment, every 26 to 38 trials. There were 10 contingency mappings for both task blocks: 2 × strongly biased (90/10; i.e. probability of reward for blue p = 0.9), 2 × moderately biased (70/30), and 2 × unbiased (50/50: as in de Berker et al., 2016). The biased mappings repeated in reverse relationships (2 × 10/90; 2 × 30/70) to ensure that over the two blocks (TB1, TB2) there were 10 stimulus-outcome contingency phases in total.

In each trial the stimuli were presented randomly to the left or right of the centre of the screen where they remained until either a response was given (left, right) or the trial expired (maximum waiting time, 2200 ms ± 200 ms). Next, the chosen image was highlighted in bright green for 1200 ms (± 200 ms) before the outcome (win, green; lose or no response, red) was shown in the middle of the screen (1200 ms ± 200 ms). At the end of each trial, the outcome was replaced by a fixation cross at an inter-trial interval of 1250 ms (± 250 ms).

Specific task instructions to participants were to select which image they predicted would reward them on each trial and to adjust their predictions according to inferred changes in the probability of reward (as in de Berker et al., 2016). All participants filled out computerised questionnaires (state anxiety STAI state scale X1, 20 items: Spielberger, 1983) and conducted practice trials as detailed in Hein et al. (2021). Critically, the state anxiety manipulation was delivered just before the first reward-learning block (TB1) to the StA group (see the following section).

### 2.3 Manipulation and assessment of state anxiety

Our StA group was instructed to complete a public speaking task in line with previous work (Feldman et al., 2004; Lang et al., 2015). This meant, as detailed in Hein et al. (2021), that StA participants were told just before TB1 that they would need to present a piece of abstract art for 5 minutes to a panel of academic experts after completing the reward-learning task, with 3 minutes preparation time. By contrast, the Cont group were informed that they would need to give a mental description of the piece of abstract artwork for the same time privately (rather than to a panel of experts, see Hein et al., 2021). Importantly, the state anxiety manipulation was then revoked in the StA group directly after completing the second reward-learning block (TB2) and before the second resting state block (R2). They were informed that the panel of experts was suddenly unavailable. Both groups, therefore, presented the artwork to themselves after completing the reward-based learning task.

To assess state anxiety, in our previous work we used the coefficient of variation (CV = standard deviation/mean) of the inter-beat intervals (IBI) as a metric of heart rate variability (HRV), as this index has been shown to drop during anxious states (Chalmers et al., 2014; Feldman et al., 2004; Gorman and Sloan, 2000; Kawachi et al., 1995; Quintana et al., 2016). Additional to this, the spectral characteristics of the IBI data were analysed to obtain an HRV proxy of state anxiety associated with autonomic modulation and parasympathetic (vagal) withdrawal (Friedman, 2007; Gorman and Sloan, 2000). HRV and high-frequency HRV (HF-HRV, 0.15–0.40 Hz) measures were derived from the R-peaks extracted from the EKG signal recorded throughout the experimental sessions (see details in Hein et al., 2021, and section **EEG acquisition and analysis** below).

In line with prior research, our previous study showed reduced HF-HRV and reduced HRV in state anxious participants relative to controls (**Figure 1C**). Reduction in these measures has been reliably shown across trait anxiety, worry, and anxiety disorders (Aikins and Craske, 2010; Friedman, 2007; Fuller, 1992; Klein et al., 1995; Miu et al., 2009; Mujica-Parodi et al., 2009; Pittig et al., 2013; Thayer et al., 1996), and thus, significant changes to these metrics suggested physiological responses consistent with state anxiety. Subjective self-reported measures of state anxiety (STAI state scale X1, 20 items: Spielberger, 1983) were taken at four points during the original Hein et al. (2021) study, but the data could not be used due to an error in STAI data collection. We showed in a separate study, however, that HRV can effectively track changes in state anxiety, as validated by concurrent changes in STAI scores (state scale; Sporn et al., 2020).

**Figure 1.**
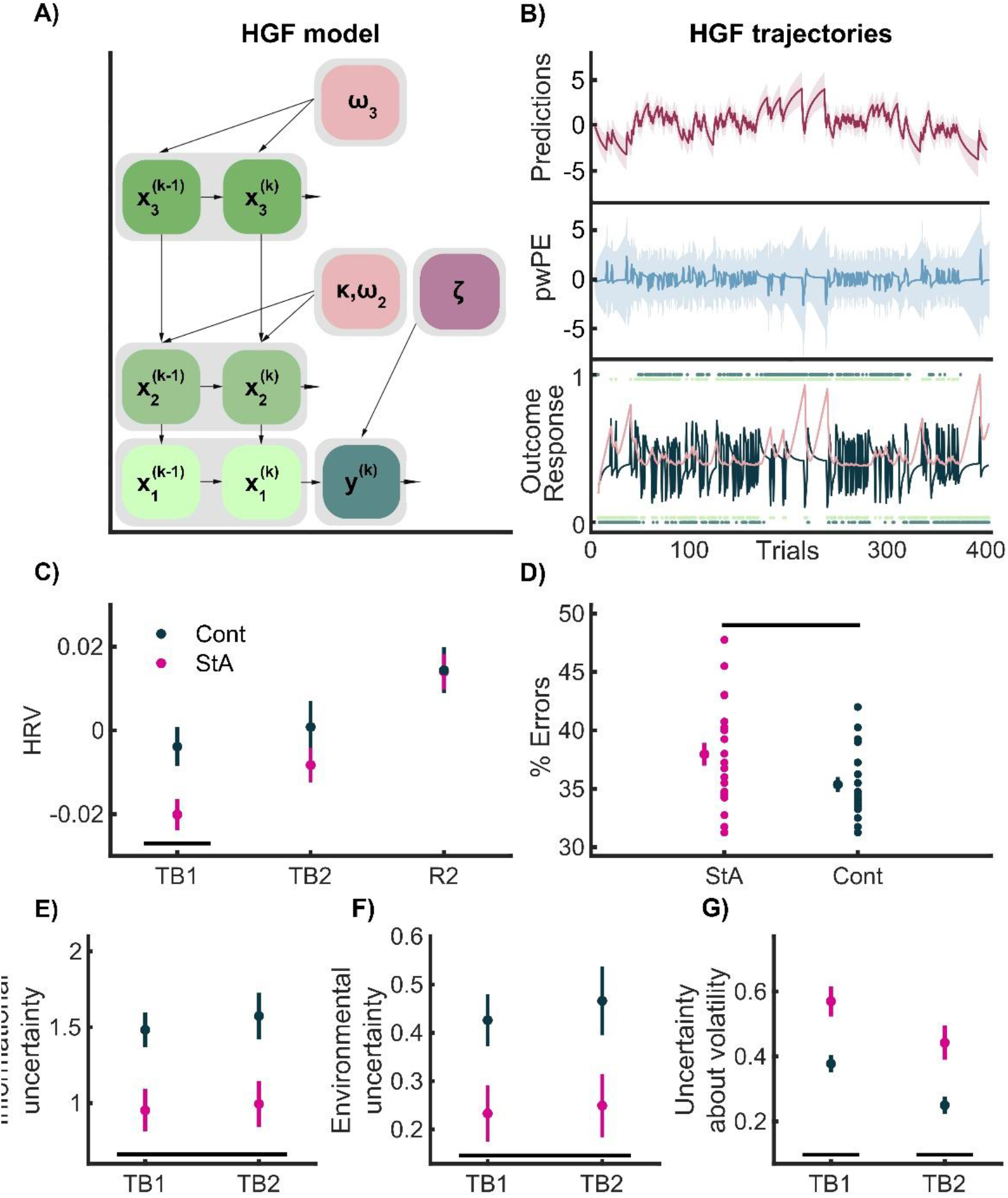
HGF model, HGF trajectory estimates and HRV, HGF and model-free results. **A)** Schematic model of 3-level HGF used in Hein et al. (2021). The free perceptual model parameters ω_2_, ω_3_ and the response parameter ζ were estimated by fitting the HGF to observed inputs (u_i_^(k)^) and individual responses (y^(k)^). **B)** HGF trajectories of the computational quantities used to form our GLM convolution regressors, from one participant. The lowest level shows the sequence of outcomes (green dots: 1 = blue win, 0 = orange win) and the participant’s responses (dark blue dots) on each trial. The black line indicates the series of prediction error (PE) responses and the pink line the precision weight. The middle layer of B) shows the trial-wise HGF estimate of pwPE about stimulus outcomes on level 2 (blue). For our GLM convolution analysis, we used unsigned values of ε_2_ as the first parametric regressor. The precision ratio included in the pwPE term, in succession, weights the influence of prediction errors on prediction updates. Predictions on level 2 are displayed on the top level (purple). We used the absolute values of predictions about the reward tendency on level 2 as the second parametric regressor. **C)** In Hein et al. (2021) a significant drop in heart rate variability (HRV, a metric of anxiety using the coefficient of variation of the inter-beat-interval of the recorded heart beats), was observed in the StA group (pink) relative to Cont (black). Panel C) shows the mean HRV (with vertical SEM bars) over the experimental task blocks 1 and 2 (TB1, TB2) and the final resting state block (R2). These blocks (TB1, TB2, R1) were normalised to the average HRV value of the first resting state block (R1: baseline). A significant effect of group and block was discovered using non-parametric 2×2 factorial tests with synchronised rearrangements. After control of the FDR at level q = 0.05, planned comparisons showed a significant between groups result (black bar) in TB1. **D)** State anxiety impeded the overall reward-based learning performance as given by the percentage of errors. In the above, the mean of each group (StA, pink, Cont, black) is provided with SEM bars extending vertically. On the right of the group mean is the individual values depicting the population dispersion. State anxiety significantly increased the error rate relative to Controls. **E)** Model-based analysis between groups revealed lower ω_2_ in StA relative to Cont produces overall lower belief estimates of *estimation* uncertainty about the reward tendency, with a main effect of the factor group (StA, pink; Cont, black). **F)** HGF results also revealed lower ω_2_ in StA leads to a main effect of group, with decreased environmental uncertainty compared with Cont **G)** State anxiety increased uncertainty about volatility (σ_3_, main effect for the factor block and group). Planned between-group comparisons also showed that StA displayed significantly higher σ_3_ relative to Cont in each task block separately (TB1, TB2, black bars).

### 2.4 Behavioural analysis and modelling

The behavioural data in our paradigm were analysed in Hein et al. (2021) using the Hierarchical Gaussian Filter (HGF, Mathys et al., 2011, 2014). This model describes hierarchically structured learning across various levels, corresponding to hidden states of the environment x_1_, x_2_,…, x_n_ and defined as coupled Gaussian random walks. Belief updating on each level is driven by PEs modulated by precision ratios, weighting the influence of precision or uncertainty in the current level and the level below. The HGF was implemented with the open-source software in TAPAS http://www.translationalneuromodeling.org/tapas).

To model learning about the tendency towards reward for blue/orange stimuli and the rate of change in that tendency (volatility), we used three alternative HGF models, and two reinforcement learning models (Hein et al., 2021). The input to the models was the series of 400 outcomes and the participant’s responses. Outcomes in trial *k* were either u^k^ = 1 if the blue image was rewarded or u^k^ = 0 if the orange image was rewarded. Trial responses were defined as y^k^ = 1 if participants chose the blue image, while y^k^ = 0 corresponded to the choice of the orange image. We tested a 3-level HGF (HGF_3_, with volatility estimated on the third level), a 2-level reduced HGF (HGF_2_, that removes the third volatility level), and a HGF where decisions are informed by trial-wise estimates of volatility (see Diaconescu et al, 2014). We additionally tested two widely used reinforcement models, a Rescorla Wagner (RW, Rescorla and Wagner, 1972) and Sutton K1 model (SK1, Sutton, 1992). Following random effects Bayesian model comparison, the model that best explained the behavioural data among participants was the 3-level HGF for binary outcomes (see **Figure 1A**). In this winning model, the first level represents the binary outcome in a trial (either blue or orange wins) and beliefs on this level feature *expected* or *irreducible* uncertainty due to the probabilistic nature of the rewarded outcome (Soltani and Izquierdo, 2019). The second level x_2_ represents the true tendency for either image (blue, orange) to be rewarding. And the third level represents the log-volatility or rate of change of reward tendencies (Bland and Schaefer, 2012; Yu and Dayan, 2005). The second and third level states, x_2_ and x_3_, are continuous variables evolving as Gaussian random walks coupled through their variance (inverse precision). Accordingly, the posterior distribution of beliefs about the two true hidden states x_2_ and x_3_ is fully determined by the sufficient statistics μ_i_ (mean, denoting participant’s expectation) and σ_i_ (variance, termed *informational* or *estimation* uncertainty for level 2; uncertainty about volatility for level 3). The coupling function between levels 2 and 3 is as follows:

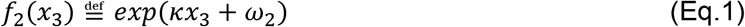

In equation (1), ω_2_ represents the invariant (tonic) portion of the log volatility of x_2_ and captures the size of each individual’s stimulus-outcome belief update independent to x_3_. Parameter ω_3_ seizes upon ‘metavolatility’: how estimates of environmental volatility evolve—with larger values articulating a belief that the changeability of the task is itself changing. Note that in our experimental task, however, the rate of change (true volatility) was constant, as the stimulus-outcome contingencies changed every 26–38 trials (similarly to de Berker et al. [2016] and Iglesias et al. [2013]). The κ parameter establishes the strength of the coupling between x_2_ and x_3_, and thus the degree to which estimated environmental volatility impacts the learning rate about the stimulus-outcome probabilities. In our implementation of the winning model, the 3-level HGF, we estimated the perceptual model parameters ω_2_, ω_3_, while k was fixed to 1. The table of prior values on the model parameters can be found in Hein et al. (2021), which includes the results of simulations carried out to assess how well the HGF_3_ estimated each free model parameter.

Paired with this perceptual model of hierarchically-related beliefs is a response model that obtains the most likely response for each trial using the belief estimates. The winning HGF model from Hein et al. (2021) used the unit-square sigmoid observation model for binary responses (; Iglesias et al., 2013; Mathys et al., 2011, 2014) and the response model parameter ζ was additionally estimated for each participant (see Table with prior values in Hein et al., 2021). We refer the reader to the original HGF methods papers for more detail on the mathematical derivations (Mathys et al., 2011, 2014), and to Hein et al. (2021) for equations included in the original results.

In the current study, we used the subject-specific trajectories of unsigned predictions and precision-weighted prediction errors on level 2 as parametric regressors for convolution GLM analysis. The arguments supporting our choice of unsigned (absolute) values for these computational quantities is given in section **Spectral Analysis** below.

The update steps for the posterior mean on level 2 and for trial k, 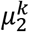, depend on the prediction error on the level below, 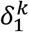, weighted by a precision ratio according to the following expression:

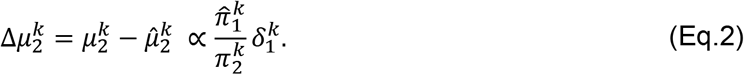

The prediction on the reward tendency before observing the outcome, 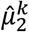, is in our winning model the expectation in the previous trial, 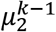. The pwPE term on level 2, on the other hand, is the product of the precision ratio and the PE about stimulus outcome: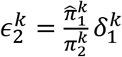. Thus, the influence of PEs on updating *μ*_2_ decreases with greater precision on that level, 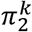, or smaller *estimation* uncertainty, 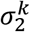. PEs contribute to larger update steps for increased precision of the prediction on the level below 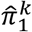. The intuition from this expression is that the less certain we are about the level we’re trying to estimate (current level), the more we should update that level using new information (prediction errors) from the level below. In addition, the more uncertain we are about the level below (reduced precision of the prediction), the less that new information should contribute to our belief updating. See Mathys et al. (2011, 2014) for detailed mathematical expressions for the HGF. Details on the free parameters of the HGF model that were estimated, including prior values, can be found in Hein et al. (2021), and are briefly mentioned in **Figure 1A**.

Trial-by-trial trajectories of the unsigned predictions 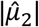 and pwPEs |ε_2_| for an exemplar participant are provided in **Figure 1B**.

### 2.5 EEG and EKG acquisition and analysis

EEG, EKG and EOG signals were recorded continuously throughout the study using the BioSemi ActiveTwo system (64 electrodes, extended international 10–20, sampling rate 512 Hz). External electrodes were placed on the left and right earlobes to use as references upon importing the EEG data in the analysis software. EKG and EOG signals were recorded using bipolar configurations. For EOG, we used two external electrodes to acquire vertical and horizontal eye-movements, one on top of the zygomatic bone by the right eye, and one between both eyes, on the glabella. For EKG we used two external electrodes in a two-lead configuration (Moody and Mark, 1982). Please refer to Hein et al. (2021) for further details on the electrophysiology acquisition.

EEG data were preprocessed in the EEGLAB toolbox (Delorme and Makeig, 2004). The continuous EEG data were first filtered using a high-pass filter at 0.5 Hz (with a hamming windowed sinc finite impulse response filter with order 3380) and notch-filtered at 48–52 Hz (filter order 846). Next, independent component analysis (ICA, runICA method) was implemented to remove artefacts related to eye blinks, saccades and heartbeats (2.3 components were removed on average [SEM 0.16]), as detailed in Hein et al. (2021).

Continuous EEG data were then segmented into epochs centred around the outcome event (win, lose, no response) from −200 to 1000 ms. Noisy data epochs defined as exceeding a threshold set to ± 100 μV were marked as artefactual (and were excluded during convolution modelling, see next section). Further to this, a stricter requirement was placed on the artefact rejection process to achieve higher quality time-frequency decomposition, as proposed for the gamma band (see Hassler et al., 2011; Keren et al., 2010). Data epochs exceeding an additional threshold set to the 75^th^ percentile+1.5·IQR (the interquartile range, summed over all channels) were marked to be rejected (Carling, 2000; Schwertman et al., 2004; Tukey, 1977). The two rejection criteria resulted in an average of 22.37 (SEM 2.4) rejected events, with a participant minimum of 80% of the total 400 events available for convolution modelling. Following preprocessing, EEG continuous data were converted to SPM 12 (http://www.fil.ion.ucl.ac.uk/spm/ version 7487) downsampled to 256 Hz and time-frequency analysis was performed (Litvak et al., 2011).

Preprocessed EEG and behavioural data files are available in the Open Science Framework Data Repository: https://osf.io/b4qkp/. All subsequent results shown here are based on these data.

### 2.6 Spectral Analysis

Prior to assessing the effect of HGF predictors on “phasic” changes in the time-frequency representations, we determined whether the average spectral power differed between state anxiety and control participants during task performance. To achieve this, we extracted the standard power spectral density (in mV^2^/Hz) of the raw data within 1–90 Hz and during task blocks TB1 and TB2 (fast Fourier transform, Welch method, Hanning window of 1 s, 75% overlap) and converted it into decibels (dB: 10*log_10_).

Standard time-frequency (TF) representations of the continuous EEG data were estimated by convolving the time series with Morlet wavelets. TF spectral power was estimated in the range 4 to 80 Hz, using a higher number of wavelet cycles for higher frequencies. For alpha (8–12 Hz) and beta (13–30 Hz) frequency ranges, we sampled the range 8–30 Hz in bins of 2 Hz, using 5– cycle wavelets shifted every sampled point (Kilner et al., 2005)—achieving a good compromise between high temporal and spectral resolution (Litvak et al., 2011; Ruiz et al., 2009). Gamma band activity (31–80 Hz) was also sampled in steps of 2 Hz, using 7-cycle wavelets.

Following the time-frequency transformation, we modelled the time series using a linear convolution model for oscillatory responses (Litvak et al., 2013). This convolution model was introduced to adapt the classical general linear model (GLM) approach of fMRI analysis to time-frequency data (Litvak et al., 2013). The main advantage of this approach is that it allows assessing the modulation of neural oscillatory responses on a trial-by-trial basis by one specific explanatory regressor while controlling for the effect of the other regressors included in the model. This control is particularly relevant in the case of stimuli or response events with variable timing on each trial. Convolution modelling of oscillatory responses has been successfully used in EEG (Litvak et al., 2013; Spitzer et al., 2016) and MEG research (Auksztulewicz et al., 2017).

In brief, the convolution GLM approach is an adaptation of the classical GLM, which aims to explain measured signals (BOLD for fMRI or time-domain EEG signals) across time as a linear combination of explanatory variables (regressors) and residual noise (Litvak et al., 2013). In convolution modelling for oscillatory responses, the measured signals are the time-frequency transformation (power or amplitude) of the continuous time series, denoted by matrix *Y* in the following expression:

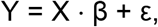

Here *Y* ∈ ℝ^t*x*f^ is defined over *t* time bins and *f* frequencies. These signals are explained by a linear combination of *n* explanatory variables or regressors in matrix *X* ∈ ℝ^t*x*n^, modulated by the regression coefficients 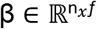. The coefficients β must be estimated for each regressor and frequency, using ordinary or weighted least squares.

The convolution modelling approach developed by Litvak et al. (2013) redefines this problem into the problem of finding time-frequency images *R_i_* for a specific type of event *i* (e.g. outcome or response event type):

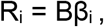

Here, *B* denotes a family of *m* basis functions (e.g. sines, cosines) used to create the regressor variables *X* by convolving the basis functions *B* with *k* input functions *U* representing the events of interest at their onset latencies, and thus *X* = *UB*. The time-frequency response images *R_i_* ∈ ℝ^p*x*f^ have dimensions *p* (peri-event interval of interest) and *f,* and are therefore interpreted as deconvolved time-frequency responses to the event types and associated parametric regressors. It is the images *R_i_* that are used for subsequent standard group-level statistical analysis. For a visual depiction of the convolution modelling of time-frequency responses, see **Figure 2**.

**Figure 2.**
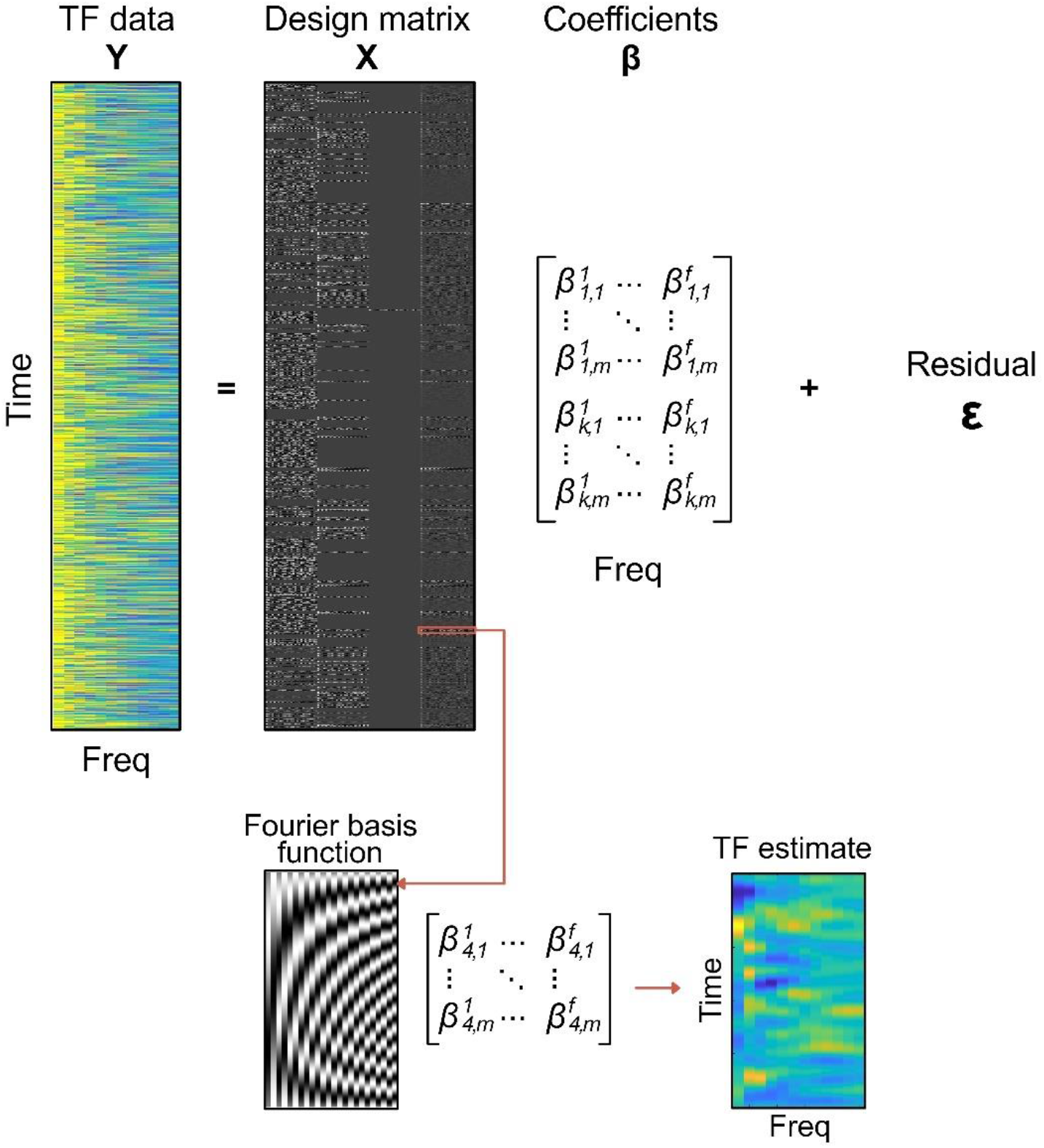
Convolution general linear model. Standard continuous time-frequency (TF) representations of the EEG signal (Y) were estimated using Morlet wavelets. In GLM, signals Y are explained by a linear combination of explanatory variables or regressors in matrix X, modulated by the regression coefficients β, and with an added noise term (ε). Our design matrix X in this example included the following regressors (columns left to right): Win, Lose, No Response, absolute pwPE on level 2, which were defined over time. Matrix X was specified as the convolution of an impulse response function, encoding the presence and value of discrete or parametric events for each regressor and time bin, and a Fourier basis function (left inset at the bottom). Solving a convolution GLM provides response images (TF estimate in the figure) that are the combination of the basis functions and the regression coefficients β_i_ for a particular regressor type i. Thus, convolution GLM effectively estimates deconvolved time-frequency responses (TF estimate, rightmost image at the bottom) to the event types and associated parametric regressors.

In our study, we were particularly interested in assessing parametric effects of computational quantities, such as pwPEs and predictions, on the time-frequency representations of the EEG data in each electrode. We implemented convolution modelling by adapting code developed by Spitzer et al. (2016) freely available at https://github.com/bernspitz/convolution-models-MEEG. The total spectral power was first converted to amplitude using a square-root transformation to conform with the GLM error assumptions (Kiebel et al., 2005; Litvak et al., 2013). Our trial-wise explanatory variables included discrete regressors coding for stimuli (blue image, orange image), responses (right, left, no response), outcome (win, lose) and relevant parametric HGF regressors: unsigned HGF model estimates of predictions on level 2 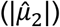 about reward outcome contingencies (hereinafter termed ‘predictions’) and precision-weighted prediction errors (pwPEs) on that level (|ε_2_|, hereinafter termed ‘pwPEs’; see **Figure 1B**). We selected the absolute value of predictions and pwPEs on level 2 because the sign in these HGF variables is arbitrary: a positive or negative value in pwPEs or predictions does not denote a win or a lose trial (see other HGF work using unsigned HGF variables as regressors, for instance, Auksztulewicz et al., 2017; Stefanics et al., 2018).

As in our previous work, pwPE on level 3 (ε_3_) updating the log-volatility estimates were excluded from this analysis due to multicollinearity: high linear correlation between pwPEs and ε_3_ (for further detail, see Hein et al., 2021). Likewise, trial-wise HGF estimates of predictions about stimulus outcomes were highly linearly correlated with predictions on the third level about volatility 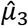(Pearson correlation coefficients ranging from −0.97 to −0.03 across all 42 participants, mean −0.7). As such, we also excluded 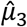 from the analysis. (For details on the impact of multicollinearity of regressors on GLMs see Mumford et al. [2015] and Vanhove [2020]). The chosen HGF pwPE and prediction regressors were consistently uncorrelated, below 0.25 in line with previous work using HGF quantities as regressors (Auksztulewicz et al., 2017; Iglesias et al., 2013; Vossel et al., 2015).

Regressor values for pwPEs were introduced at the latency of the outcome regressor and thus allowed us to assess the parametric effect of pwPEs about stimulus outcomes on the time-frequency responses in a relevant peri-event time interval. Although previous work analysed the effect of pwPEs on neural responses up to 1000 ms, we showed in Sporn et al. (2020) that pwPEs during reward learning can modulate neural oscillatory responses in the beta band up to 1600 ms, and these responses are dissociated between anxiety and control groups. The recent studies by Bauer et al. (2014) and Palmer et al. (2019) also showed that the latency of PE and pwPE effects on neural activity can extend up to 2 seconds. Accordingly, the pwPE convolution model was estimated using a window from −200 to 2000 ms relative to the outcome event, and the statistical analysis focused on the 100-1600 ms interval (see next section).

Concerning the prediction regressor, we considered different time intervals in which we could capture neural oscillatory responses to predictions. This is a challenging task acknowledged before (Diaconescu et al., 2017), as the neural representation of predictions likely evolves gradually from the outcome on the previous trial to the outcome on the current trial. It is thus not expected to be locked to a specific event. This explains why most of the previous work using the HGF framework excluded predictions as a regressor for GLM analysis. Here we followed Auksztulewicz et al. (2017), who analysed predictions locked to the cue, and Palmer et al. (2019), who assessed a wide interval surrounding the movement (response); note that in the Palmer et al. (2009) study, the motor response was the last event in each trial (i.e. there was no additional response feedback). We thus hypothesised that the neural representation of predictions on the reward outcome contingencies could be captured by focusing on two complementary windows of analysis: (i) an interval following the stimulus presentation (stimulus-locked); (ii) an interval preceding the outcome on the current trial (outcome-locked). Unlike the targeted pwPE analysis described above, the analysis of the prediction regressor was exploratory as we did not have a strong hypothesis regarding which of both time windows would preferentially reflect prediction-related neural modulations.

To assess the stimulus-locked parametric effect of predictions on the time-frequency responses, we run a convolution GLM in a time interval from −200 to 2000 ms. For the outcome-locked parametric effect of predictions, the convolution GLM was run from −2500 to 0 ms. This later interval extended to −2500 to allow for the presence of a baseline interval in every trial prior to the preceding stimulus. Thus, two separate convolution GLMs were run with the prediction regressor modulating neural activity locked to either the stimulus or outcome events. These broad windows were further refined in our statistical analysis (see next section).

In all alpha-beta convolution GLM analyses, discrete and parametric regressors were convolved with a 12th-order Fourier basis set (24 basis functions, 12 sines and 12 cosines), as in Litvak et al. (2013). For convolution models run from −200 to 2000 ms locked to an event type, using a 12th-order basis functions set allowed the GLM to resolve modulations in the TF responses up to ~ 5.5 Hz (12 cycles / 2.2 seconds; or 183 ms). For the outcome-locked GLM run from −2500 to 0 ms, the 12th-order Fourier basis set resolves frequencies up to ~5 Hz. Our choice of a 12th order set was compatible with the temporal extent of the pwPE and prediction effects on alpha-beta oscillatory activity reported in previous work (200-400 ms-long effects in (Auksztulewicz et al., 2017) up to 2000 ms-long effects in Palmer et al. (2019). In the case of gamma oscillations modulating pwPEs, we considered a higher order basis function set to allow for potentially faster gamma effects to be resolved. Using a 20th-order Fourier basis set on the gamma-band convolution GLM within −200 to 2000 ms enabled resolving modulations in the TF responses up to ~ 9 Hz (20 cycles / 2.2 seconds; or 110 ms).

### 2.7 Statistical analysis

The time-frequency images (in arbitrary units, a.u.) from the convolution model were subsequently converted to data structures compatible with the FieldTrip Toolbox for statistical analysis (Oostenveld et al., 2011). We used permutation tests with a cluster-based threshold correction to control the family-wise error rate (FWER) at level 0.05 (5000 iterations; Maris and Oostenveld, 2007; Oostenveld et al., 2011). These analyses were conducted with spatio-spectral-temporal data, after averaging the time-frequency responses within each frequency band (alpha, beta and gamma ranges). Thus, we run the cluster-based permutation tests along the spatial (64 channels), frequency-band (3) and temporal dimensions (FWER-controlled). The statistics approach consisted of investigating separately within and between-group effects. The within-group level analysis used dependent-samples two-sided tests and aimed to assess whether the neural oscillatory responses to the HGF regressors were larger or smaller than baseline levels. Next, we separately evaluated between-group effects of HGF regressors on oscillatory responses using one-sided tests (N = 21 Cont, 21 StA). This allowed us to test our hypothesis of increased alpha and beta activity and reduced gamma activity in StA compared to Cont. In the case of two-sided tests, the cluster-based test statistic used as threshold the 2.5-th and the 97.5-th quantiles of the t-distribution, whereas we used the 95th quantile of the permutation distribution as critical value in one-sided tests.

#### Analysis of the pwPE regressor

At the within-group level, we assessed the changes in time-frequency activity relative to a baseline period (given independently below) separately in StA and Cont groups (N = 21 each). Note that in convolution modelling for oscillatory responses the TF images are not baseline corrected as in standard approaches (no subtraction or division by the average baseline level). Instead, the baseline activity is estimated similarly to the post-event activity taking into account the latency variation of different events in the continuous recording (Litvak et al., 2013). Thus, TF images are not centred at 0 amplitude during the baseline period.

For the within-group analysis of the pwPE regressor, we contrasted the time-frequency images between an interval from 100 to 1600 ms post-outcome and a baseline level averaged from −200 to 0 ms, separately in each group. The 100–1600 ms time window of analysis encompasses the effects from our previous single-trial ERP study (Hein et al., 2021) and our work on the modulation of beta oscillatory responses by pwPEs during motor learning in state anxiety, which revealed effects between 400-1600 ms (Sporn et al., 2020). Between-group differences in TF representations of pwPE were separately assessed. This analysis was also conducted within 100-1600 ms. Overall, we controlled the FWER at level 0.05 to deal with the issue of multiple comparisons emerging from the spatial (64) × spectral (3) × temporal dimensions.

#### Analysis of the predictions regressor

Within-group level statistical analysis of the stimulus-locked time-frequency images of the predictions regressor focused on the range 100–1000 ms, and relative to an average pre-stimulus baseline level from −200 to 0 ms. This target window for statistical analysis balanced the evidence from previous work (Auksztulewicz et al., 2017; Palmer et al., 2019). In the current study, participants’ response preparation and execution times fell within the interval 100–1000 ms (reaction time was 598 ms on average, SEM 130 ms). However, the effects of the response were factored out from the prediction-related oscillatory activity by including the response regressor in the convolution GLM. This was validated in a control analysis that assessed the effect of the response regressor in the same time window, between 100–1000 ms stimulus-locked, to confirm independent changes in sensorimotor regions.

Within-group statistical analysis of the outcome-locked effects of predictions was conducted in a similar window 100–1000 ms preceding the outcome event (that is, from −1000 to −100 ms before the outcome). Activation in this interval was contrasted to a baseline level of 200 ms, from −2300 to −2100 ms. This baseline period was calculated to safely precede stimuli presentation across all trials, during which participants were fixating on a central point on the monitor. As mentioned above, to confirm independent changes in sensorimotor regions in response to the response regressor, we used an identical window in an additional control analysis.

The between-group level stimulus-locked analysis of predictions was conducted within 100–1000 ms. The outcome-locked analysis targeted the interval from −1000 to −100 ms, as mentioned above. As mentioned above, the FWER was controlled at level 0.05.

## 3. Results

### 3.1 Previous results: Biases of state anxiety on processing uncertainty

In Hein et al. (2021) we showed state anxiety (StA) significantly reduced HRV and HF-HRV (0.15–0.40 Hz) relative to the control group (Cont, **Figure 1C**). This outcome suggested that our state anxiety manipulation had successfully modulated physiological responses in a manner consistent with changes in state anxiety (Friedman, 2007; Fuller, 1992; Klein et al., 1995; Miu et al., 2009; Pittig et al., 2013). We further showed that state anxiety significantly increased the percentage of errors made during reward learning when compared to the control group (**Figure 1D**). In parallel to the cardiovascular and behavioural changes induced by the anxiety manipulation, by modelling decisions with the HGF, we found that state anxiety impaired learning through lowering the perceptual model parameter ω_2_. Smaller ω_2_ values reflect that StA learners updated their beliefs about reward outcomes in smaller steps than controls. In addition, lower ω_2_ values in StA were further related to the altered processing of three forms of uncertainty. First, we found significantly reduced *estimation* (belief) uncertainty (σ_2_) about the reward tendency in StA relative to Cont (**Figure 1E**). This bias in StA indicates that new information has a smaller impact on the update equations for beliefs about the tendency of reward (level 2). State anxious individuals also exhibited an underestimation of environmental uncertainty when compared with controls (**Figure 1F**). However, volatility uncertainty (σ_3_) increased in StA relative to Cont (**Figure 1G**). These HGF model-based results were aligned with the results of our separate model-free behavioural analysis as mentioned above, demonstrating a significantly higher error rate in StA during reward-learning performance (**Figure 1D**).

### 3.2 Time-frequency responses

#### 3.2.1 General modulation of spectral power

The average raw spectral power during task performance did not differ between state anxiety and control participants (P > 0.05, cluster-based permutation test; **Supplementary Figure 1**). This finding suggested that the state anxiety manipulation did not significantly modulate the general spectral profile of oscillatory activity during task performance, as we showed in a recent study (Sporn et al., 2020).

#### 3.2.2 Precision-weighted prediction errors about stimulus outcomes

The overall time course of the parametric modulation of alpha (8–12 Hz) and beta (13-30 Hz) oscillatory activity by pwPEs about stimulus outcomes is displayed in **Figure 3A** and **Figure 4A**, respectively. On the within-subject level, there was a significant decrease relative to baseline in alpha and beta activity in the control group (one negative cluster, P = 0.0002, two-sided test, FWER-controlled. Effect within 600-1400 ms for alpha, and 400–1120 ms for beta activity). No significant clusters were found in the gamma band. The alpha-band effect originated in central parietal electrodes and later spread across the whole scalp (**Figure 3B**), while the beta-band modulation had a widespread topography and started earlier than the alpha-band effect (at 400 ms; **Figure 4B**). In the StA group, a negative cluster was also found, corresponding to a decrease from baseline in alpha and beta activity (P = 0.0054, two-sided test; 600–1000 ms for alpha, 440–1000 ms for beta). The StA alpha-band effect also emerged in centro-parietal electrodes but later shifted to frontocentral electrodes (**Figure 3C**). In the beta range, the negative modulation of oscillatory activity in StA had a right frontocentral and left centro-parietal distribution (**Figure 4C**).

**Figure 3.**
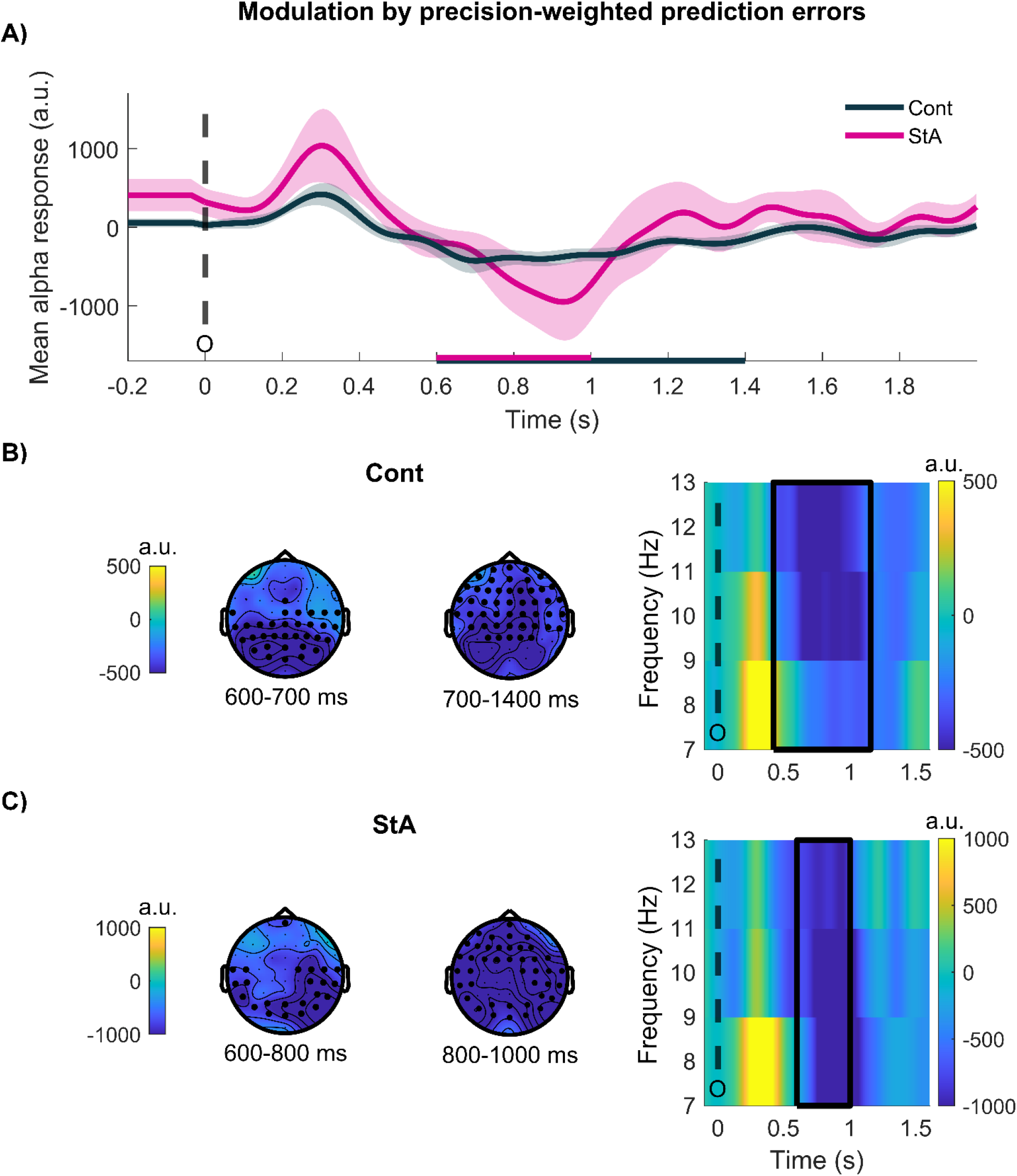
Alpha activity is modulated by precision-weighted prediction errors about stimulus outcomes: within-group effects. **A)** Time course of the average alpha response (8–12 Hz) to pwPEs in each group (Controls, black; StA, pink), given in arbitrary units (a.u). The time intervals correspond to the dependent-samples significant clusters (B, C) in each group, and are denoted by horizontal bars on the x-axis. **B)** Electrode-level correlates of pwPEs in alpha activity in the Cont group. One negative cluster was found spanning the alpha and beta frequency range. The within-group effect in alpha activity emerged between 600– 1400 ms (P = 0.0002). Left: The topographic distribution of this effect starts in posterior centroparietal regions and expands across all electrode regions. Right: Time-frequency images for pwPE on level 2, averaged across the cluster electrodes. The black dashed line marks the onset of the outcome, and black squares indicate the time-frequency range of the significant cluster. **C)** Same as (B) but in the StA group. We found a significant negative cluster also in the alpha and beta-band ranges. The alpha-band effect was found between 600–1000 ms (P = 0.0054), starting in posterior central electrodes and spreading to frontocentral electrodes later. Dashed and continuous black lines denote outcome onset and the extension of the significant cluster in the time-frequency range, as in (B).

**Figure 4.**
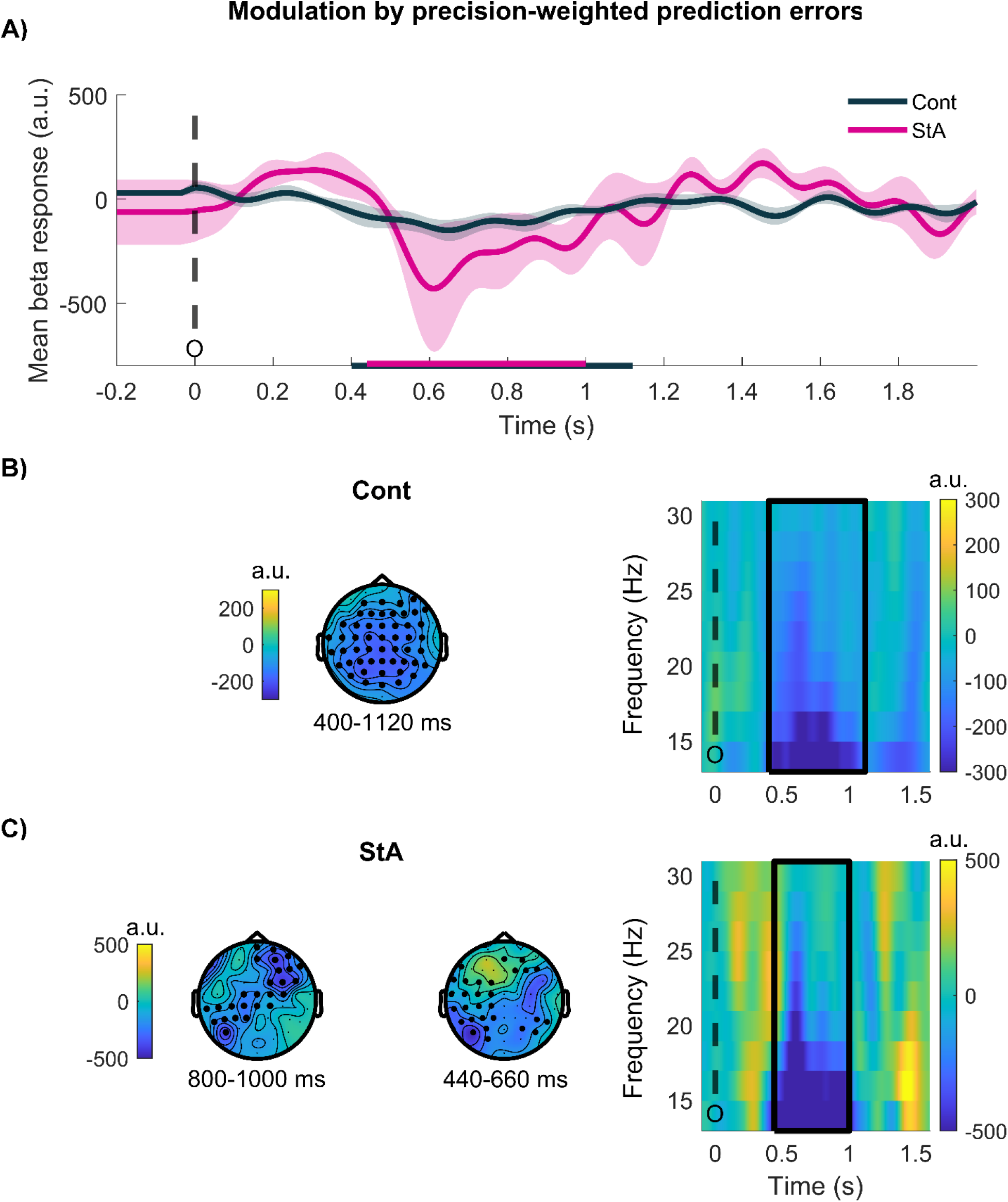
Beta activity is modulated by precision-weighted prediction errors about stimulus outcomes: within-group effects. **A)** Time course of the average beta response (12–30 Hz) to pwPEs in each group (Controls, black; StA, pink), given in arbitrary units (a.u)). The time intervals corresponding to the dependent-samples significant clusters (B, C) in each group are denoted by horizontal bars on the x-axis. **B)** Electrode-level correlates of pwPEs in beta activity in the Cont group. The significant negative cluster found in the alpha and beta ranges, had a temporal distribution between 400–1120 ms in the case of beta activity (P = 0.0002, FWER-controlled due to multiple comparisons arising from testing across space x frequency-band x time dimensions). Left: The topographic distribution of the beta-band effect is across the entire scalp. Right: Time-frequency images for pwPE on level 2, averaged across the significant cluster electrodes. The black dashed line marks the onset of the outcome, and black squares indicate the time-frequency range of the significant cluster. **C)** Same as (B) but in the StA group. As reported in Figure 3, we found a significant negative cluster across the alpha and beta band, with a latency of 440–1000 ms for beta activity (P = 0.0054, FWER-controlled). The beta modulation started in posterior central electrodes and later spread to frontocentral electrodes. Dashed and continuous black lines denote outcome onset and the extension of the significant cluster in the time-frequency range, as in (B).

Complementing the within-subject results, independent-samples statistical tests across the alpha, beta and gamma ranges revealed one significant positive cluster in the beta range (between 1200–1570 ms, P = 0.027, one-sided test; FWER-controlled). This effect was associated with higher beta activity at left sensorimotor and frontocentral electrodes in StA relative to Cont (**Figure 5AB)**. The individual average of beta-band activity in the significant cluster is shown in **Figure 5C.** Of note, in StA, a qualitative comparison of the sensorimotor and frontocentral beta activity associated with the significant cluster of the between-group statistical analysis revealed a greater activity increase in the sensorimotor than in the frontocentral electrode region (**Figure 5D**). In the control group, the beta response to pwPE decreased in both electrode regions, but the reduction was more pronounced in frontocentral electrodes (**Figure 5E**). There were no additional significant clusters associated with between-group differences in the alpha or gamma ranges (see illustration of gamma responses to pwPE in **Supplementary Figure 2**).

**Figure 5.**
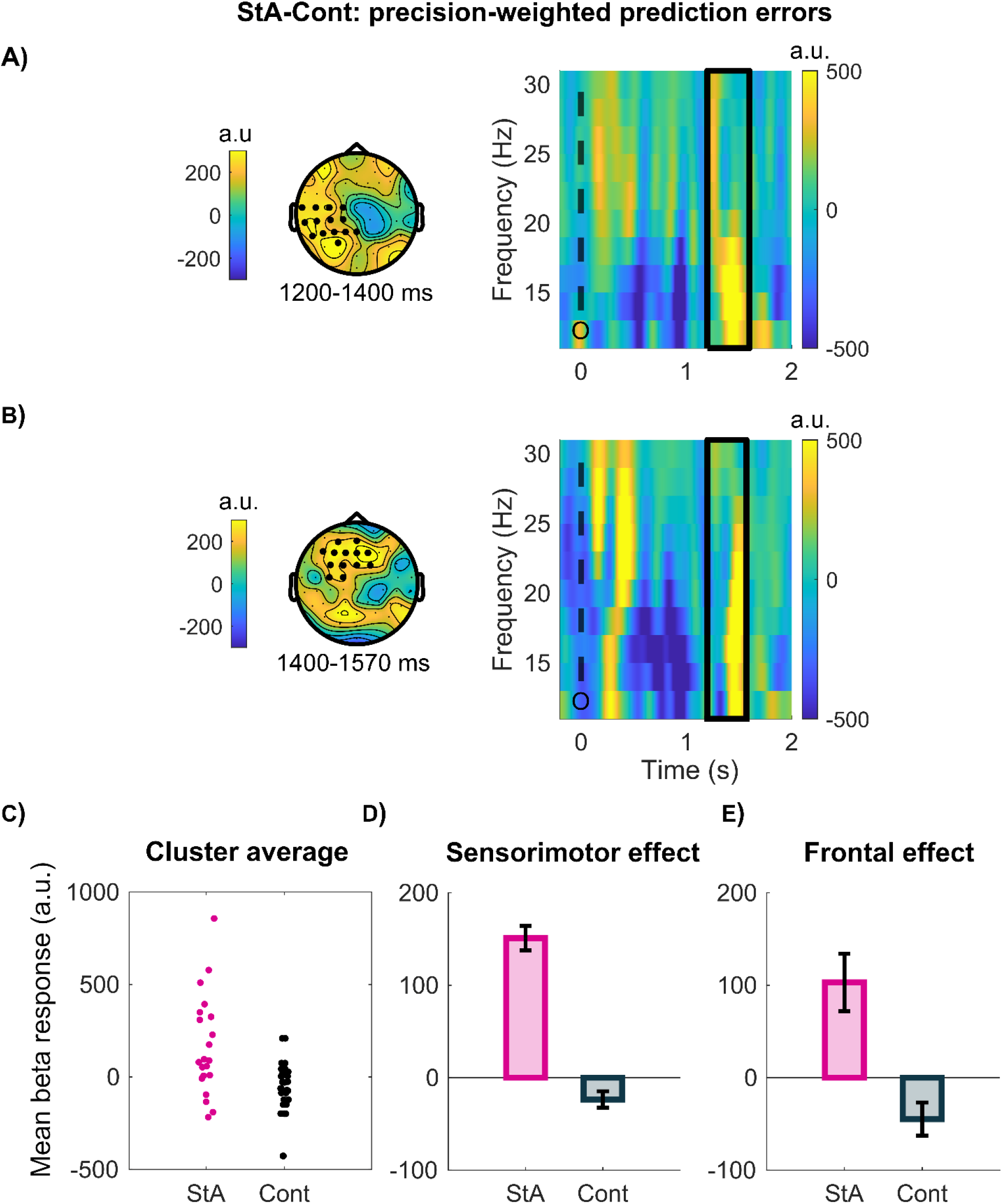
Between-group effects of pwPEs on beta oscillatory activity. **A-B)** Between-group differences in the oscillatory activity (alpha 8-12 Hz, beta 13-30 Hz, and gamma 31-90 Hz) modulations by pwPEs about stimulus outcomes were found exclusively in the beta band (one significant positive cluster between 1200–1570 ms; P = 0.0270), FWER-controlled). **A)** The topographic distribution of this effect starts early in left posterior parietal regions and **B)** later shifts to frontocentral electrodes. The time-frequency images on the right panels correspond to the electrode selection on the left topographic panels and are given in arbitrary units (a. u.). Note that there was one single significant cluster represented by the solid black rectangle; dashed black line ‘O’ represents the time of the outcome. **C)** The average beta response (a. u.) to pwPE for individuals in each group (StA, pink; Cont, Black) in the cluster. **D)** The modulation of time-frequency responses to pwPEs in beta displayed separately in sensorimotor (left) and frontal (right) electrodes pertaining to the significant cluster. Pink bars represent results in the state anxiety group (StA), whereas the control group (Cont) is denoted by black bars. Black “error” bars indicate the standard error of the mean (SEM).

### 3.2.3 Predictions about reward tendency

#### Stimulus Locked

When assessing within-group level modulations in oscillatory activity by the prediction regressor, there were no significant effects, neither in the Cont or StA group (P > 0.05, FWER-controlled). Between-group statistical analysis revealed that predictions about the reward tendency are associated with significantly higher levels of beta activity in StA than in Cont across frontocentral and parietal electrodes (one positive cluster in the beta band only, from 200 to 640 ms, P = 0.04, one-sided test, FWER-controlled; **Figure 6AB**). There were no additional significant clusters extending to the alpha range.

**Figure 6.**
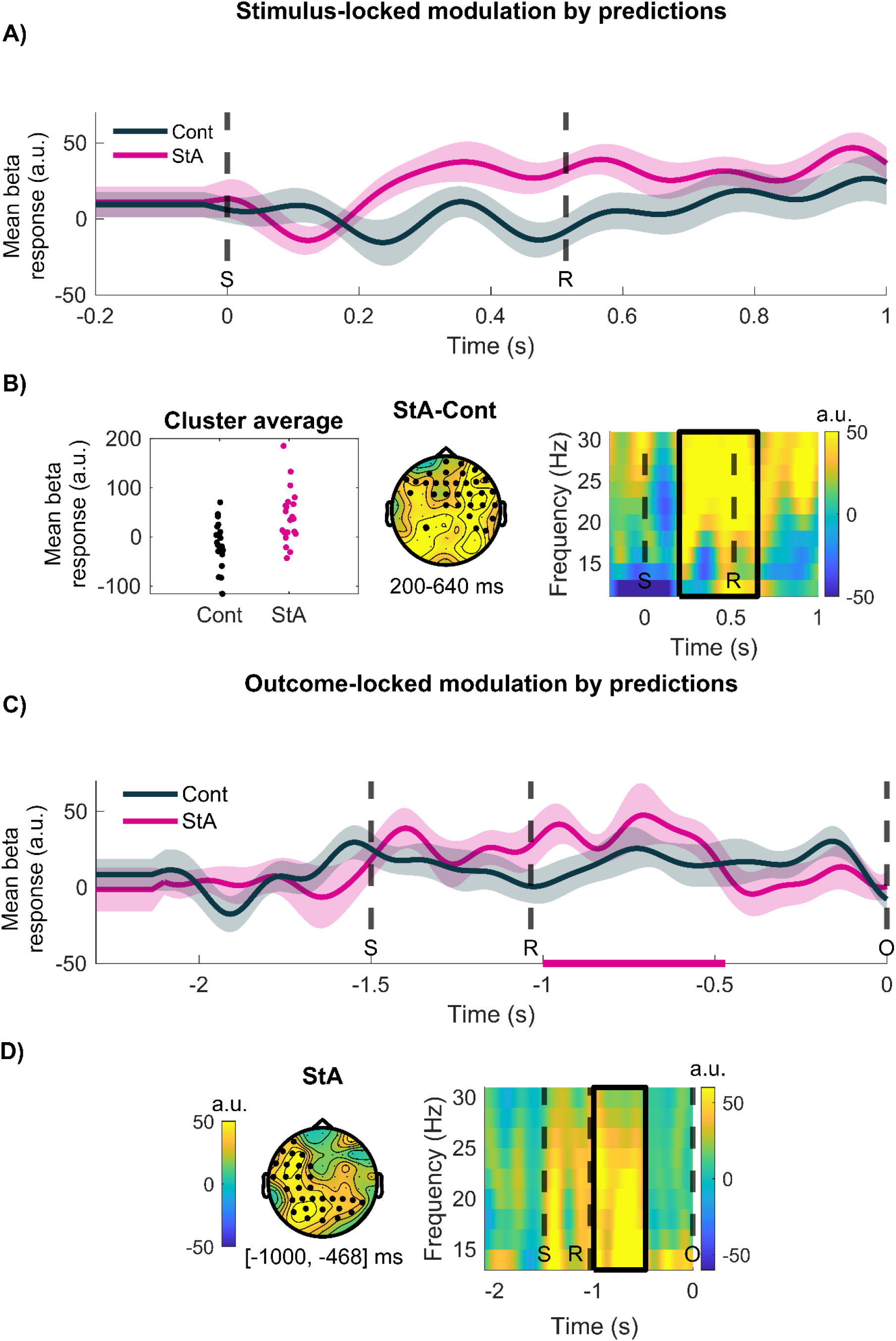
Stimulus-locked modulation of beta activity by predictions. **A)** Average time course of stimulus-locked beta activity modulated by predictions in Cont (black) and StA (pink). Modulation of time-frequency images by a regressor is dimensionless, and thus given in arbitrary units (a.u). **B)** In the leftmost column is the average beta activity (a. u.) in the cluster to pwPE for individuals in each group (StA, pink; Cont, Black). An independent-samples test on beta activity revealed a significant increase in StA relative to Cont from 200 to 640 ms across parietal and frontocentral electrodes (one significant positive cluster, P = 0.04, FWER-controlled). **C)** Outcome-locked modulation of beta activity by predictions. Average time course in a.u. of outcome-locked beta activity reflecting modulation by predictions in Cont (black) and StA (pink). The significant within-group effect in StA is shown by the pink horizontal bar on the x-axis. **D)** Within-group statistical analysis with dependent-samples cluster-based permutation tests revealed one positive cluster in the state anxious group ([−1000, −468] ms, P = 0.0106, two-sided test, FWER-controlled), reflecting increased beta activity in centroparietal and left frontal electrodes during processing predictions. Solid black lines represent the time and frequency of the significant cluster. Dashed black lines represent the average time of the stimuli presentation ‘S’, participant’s response ‘R’ and the outcome ‘O’.

The between-group effect of predictions on beta activity was not confounded by any concomitant effect of motor responses on the neural oscillatory responses, as we had included a response regressor in this analysis. A control analysis on this independent-samples effect of the response regressor on beta activity showed no significant difference between the two groups (see **Supplementary Figure 3A**).

#### Outcome-locked

**Figure 6C** displays the time course of the parametric effects of predictions on outcome-locked beta activity. State anxious participants exhibited a significant increase from baseline in beta oscillatory activity (one significant positive cluster from −1000 to −468 ms, P = 0.0106, two-sided test, FWER-controlled). This effect peaked at central parietal and left frontocentral electrodes (see **Figure 6D**). There were no significant changes from baseline in alpha or beta oscillatory activity for the control group participants. Neither did we find significant between-group differences in outcome-locked alpha or beta activity. Like in our stimulus-locked results, the significant outcome-locked increase from baseline in beta oscillatory activity in the StA group was not confounded by motor modulation, as this was included as a separate regressor in the convolution model. A control analysis of the effect of the response regressor on beta activity yielded non-significant changes from baseline in StA (**Supplementary Figure 3B**).

## 4 Discussion

This study investigated how anxiety states modulate the oscillatory correlates of low-level reward predictions and prediction errors during learning in a volatile environment. Because in generative models of the external world precision weights regulate the influence that PEs have on updating predictions (Feldman and Friston, 2010; Friston, 2010), we assessed the neural oscillatory responses to precision-weighted PEs (pwPEs), rather than PEs alone, similarly to Auksztulewicz et al. (2017). We tested this by re-analysing data from our previous study, which investigated Bayesian predictive coding (PC) in state anxiety and showed that anxious individuals overestimate how precise their belief about the reward tendency is, attenuating pwPEs on that level and decreasing learning (Hein et al., 2021). In the current study, trial-wise model estimates of predictions and pwPEs were used as parametric regressors in a convolution model to explain modulations in the amplitude of oscillatory EEG activity (Litvak et al., 2013).

Consistent with our hypotheses, we found that state anxiety alters the spectral correlates of prediction and pwPE signalling. Anxiety states amplified beta oscillations during processing of predictions about the reward tendency. This outcome may represent a stronger reliance on prior beliefs (Bauer et al., 2014; Sedley et al., 2016), downweighting the role of PEs in updating predictions and suppressing gamma responses (Bauer et al., 2014). While pwPEs did not significantly modulate gamma activity as a function of anxiety, we found increased beta modulations during the processing of pwPEs in state anxiety. This outcome is aligned with our recent findings in anxiety during reward-based motor learning (Sporn et al., 2020). In addition, as detailed below, the result can be reconciled with hypotheses from generalised PC (Brown and Friston, 2013; Feldman and Friston, 2010) in which attention modulates precision weights on PEs through changes in synaptic gains and lower frequency oscillations (Bauer et al., 2014; Sedley et al., 2016). Overall, our results extend computational work on maladaptive learning in anxiety, suggesting that altered beta frequency oscillations may explain impeded reward learning in anxiety, particularly in volatile environments (Browning et al., 2015; Piray et al., 2019; Pulcu and Browning, 2019).

### Oscillatory correlates of precision-weighted prediction errors in state anxiety

In Hein et al. (2021), a 3-level HGF model best described learning behaviour. Key findings were that state anxiety decreased the overall learning rate and led to an underestimation of environmental and *estimation* uncertainty about the reward tendency. As lower *estimation* uncertainty (inverse precision) about stimulus outcomes drove smaller pwPEs on that level, decreasing learning rates, here we predicted lower gamma activity during processing pwPEs in the state anxiety group. Given that both enhanced gamma and suppressed beta (and alpha) activity have recently been associated with pwPE during perceptual learning (Auksztulewicz et al. 2017) and with processing unexpected stimuli (Bastos et al., 2020), we also hypothesised concurrent higher alpha and beta modulation in state anxiety during pwPE signalling. More generally, gamma oscillations are anticorrelated with beta (and alpha) oscillations across the cortex, as shown for sensorimotor processing and working memory (Hoogenboom et al., 2006; Lundqvist et al., 2016, 2018, 2020; Miller et al., 2018; Potes et al., 2014).

Our results provide novel insight into how rhythm-based formulations of (Bayesian) PC—initially proposed for sensory processing—can be extended to reward-based learning. Our findings show that unsigned pwPEs about the stimulus outcomes first decreased both alpha and beta activity 400–1000 ms post-outcome, separately in each group, suggesting that attenuation of lower frequency responses is associated with processing pwPEs independently of anxiety. Subsequently, during 1200–1570 ms, state anxiety relative to controls increased beta responses in sensorimotor and frontocentral electrode regions. This latter effect is closely aligned with the effects of state anxiety on beta activity (power and burst events) during processing pwPEs for reward-based motor learning (Sporn et al., 2020). Reduced alpha-beta activity was also linked to pwPEs in Auksztulewicz et al. (2017), yet this effect was paralleled by increased gamma oscillatory activity. We failed to find any effects of pwPEs on gamma activity, limiting the interpretation of the results. However, different accounts outlined below could partially explain our lack of gamma-band effects.

Suppression of PEs conveyed by gamma oscillations can occur through two main mechanisms: (1) the inhibitory effects of top-down predictions, and (2) postsynaptic gain regulation (Bauer et al., 2014; Brown and Friston, 2013; Larkum et al., 2004). Both mechanisms could partly account for our findings. On the one hand, the greater beta activity associated with predictions in state anxiety would convey inhibitory input to superficial pyramidal neurons encoding PEs, decreasing gamma (Bastos et al., 2012; Sedley et al., 2016). On the other hand, in state anxiety, the lower *estimation* uncertainty σ_2_ reduces the precision ratio modulating PEs: 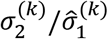 or 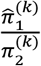 (**Eq. 2**). Accordingly, pwPEs and associated gamma activity would decline.

Mechanistically, precision is thought encoded via postsynaptic gain, modulated by neurotransmitters and attentional processes (Bauer et al., 2014; Feldman and Friston, 2010; Friston and Kiebel, 2009; Moran et al., 2013). Empirical investigations of sensory PEs link alpha and beta oscillations to the encoding of the precision of predictions (Bauer et al., 2014; Palmer et al., 2019; Sedley et al., 2016), which is the numerator of the precision ratio 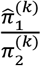 in the HGF update equations on level 2, and not affected by anxiety our study. In visuomotor adaptation tasks, high sensory uncertainty after visual perturbations (low 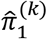) is followed by greater post-movement beta power, reflecting a greater reliance on priors (Palmer et al., 2019) and increased confidence in the feedforward estimations relative to sensory feedback (Tan et al., 2016).

Because we investigated reward-based learning to capture the effects of external motivational feedback on learning biases in anxiety, the relevant precision term in our computational model was π_2_, the precision of the posterior belief in the reward tendency. Increased precision π_2_, as we observed in state anxiety, could explain the increased beta activity in this group during encoding pwPEs, reflecting smaller updates to predictions and diminishing the magnitude of gamma activity. These accounts don’t explain, however, the lack of gamma results on the within-subject level. Our follow-up work with MEG will aim to elucidate the role of gamma oscillations during belief updating in healthy and subclinical anxiety samples.

A further consideration for future work is the role of dopamine in mediating anxiety-related changes in alpha-beta activity during Bayesian PC. Dopamine is considered a relevant signal encoding the precision of beliefs during reward learning (FitzGerald et al., 2015) and is associated with sensorimotor beta oscillations in research on Parkinson’s (Haumesser et al., 2021; Jenkinson and Brown 2011). Dopamine also plays an important modulatory role in fear and anxiety disorders (de la Mora et al., 2010). Pharmacological interventions are a powerful approach to link computational quantities to neurotransmitters (Marshall et al., 2016; Vossel et al., 2014), and thus may offer insights into the process of biased learning in anxiety.

More generally, EEG/MEG studies consistently show that frontocentral beta oscillations are modulated by positive reward feedback or predicting cues (Bunzeck et al., 2011; Cunillera et al., 2012; Marco-Pallares et al., 2008;). These effects seem to stem from cortical structures linked to the reward-related fronto-subcortical network, such as the PFC (HajiHosseini et al., 2012; Mas-Herrero et al., 2015; O’Doherty, 2004). These studies, however, did not directly model the update of reward predictions via PEs. Beyond the Bayesian PC interpretations, a common view is that reduced beta activity in the prefrontal, somatosensory, and sensorimotor territories facilitates the encoding of relevant information to shape ongoing task performance (Engel and Fries, 2010; Schmidt et al., 2019, Shin et al., 2017). Accordingly, state anxiety could be more broadly associated with disrupting processing of relevant information through changes in beta oscillations, in line with some of the evidence on EEG markers of social anxiety disorders (Al-Ezzi et al., 2020).

While anxiety commonly disrupts performance in higher-order executive tasks, such as during decision making and probabilistic learning (Grupe and Nitschke 2013; Miu et al. 2008; Remmers and Zander 2018), adaptive benefits are frequently reported from hypervigilant states when performing perceptual or sensory processing tasks, including threat detection (Bach, 2015; Cornwell et al., 2017; Grillon, 2008; Robinson et al., 2011). In pathological anxiety, this hypervigilance to potential aversive stimuli is exaggerated and overapplied to everyday unthreatening contexts. Given this discrepancy between anxious responses for perceptual and executive tasks, we speculate that the neural oscillatory patterns reported here might be reversed for anxious individuals during learning about aversive or threatening stimuli. One might expect in those learning scenarios that anxiety will amplify gamma frequency responses at the expense of beta frequency activity relative to controls. This represents an important future direction for experimental research in anxiety; however, one challenge is to disentangle the effects of different anxiety induction methods on these neural and behavioural responses and how these are distinct from clinical disorders (Grillon, 2019). By identifying and explaining these learning difficulties and differences processing uncertainty in anxiety, research implications could inform treatment and learning-based therapies (Moutoussis et al. 2018).

### Biased predictions in state anxiety are associated with enhanced beta oscillations

Capturing neural modulations by predictions is challenging (Diaconescu et al., 2017). The neural representation of predictions could develop anywhere between the previous and current trial’s outcome. To address this, we separately analysed oscillatory correlates of predictions post-stimulus and pre-outcome. Between-group effects were obtained exclusively in the stimulus-locked analysis, corresponding with an increase in beta activity between 200–640 ms in the StA relative to the control group, with a widespread topography. This effect was paralleled by a significant beta activity increase in the state anxiety group, yet exclusively in the outcome-locked representation, from −1000 to −500 ms prior to the outcome. This effect extended across central, parietal, and frontal electrode regions. Our analysis focusing on two different yet dependent windows was exploratory; we did not have a strong hypothesis concerning which time interval would be best suited to assess the effect of anxiety on neural oscillatory correlates of predictions, given the gradual modulation of predictions argued before (Diaconescu et al., 2017). Future studies will need to clarify this aspect, but between-group comparisons likely benefit from a stimulus-locked approach when assessing the oscillatory correlates of predictions.

Previous work consistently linked alpha and beta oscillatory power to encoding predictions— potentially down-modulating precision weights (Auksztulewicz et al., 2017; Bauer et al., 2014; Sedley et al., 2016). Extant work, however, focused on sensory predictions and healthy control participants, which leaves open the question of how aberrant affective states may interact with oscillatory correlates of prediction signals. In our study, interpretation of results in healthy controls is limited given the lack of a significant modulation by prediction in this group. EEG predominantly samples brain electrical activity in superficial cortical layers (Buzsáki et al., 2012; Lopes da Silva, 2013) and may be less sensitive to oscillatory activity associated with predictions in deep layers, at least in a normal physiological state. In temporary anxiety, by contrast, abnormally precise posterior beliefs would suppress PEs, strengthening predictions and exacerbating the associated beta activity. Further investigation is needed to identify the oscillatory responses to reward-based prediction and PE signalling in healthy controls, opening up rhythm-based accounts of Bayesian PC to reward learning. Above all, our findings extend recent computational work on learning difficulties in anxiety (Browning et al., 2015; de Visser et al., 2010; Huang et al., 2017; Lamba et al., 2020; Miu et al., 2008; Piray et al., 2019). We propose enhanced beta oscillations as one relevant neurophysiological marker associated with state anxiety’s impairment of reward learning; these may maintain overly precise unyielding feedforward estimations of reward resistant to revision that inhibit PE encoding.

## Acknowledgements

Funding: This study was supported by Goldsmiths University London, funded by the Economic and Social Research Council (ESRC) and the South East Network for Social Sciences (SeNSS) through grant ES/P00072X/1, and the Basic Research Program at the National Research University Higher School of Economics (Russian Federation).

## Author contributions

M.H.R. and T.P.H. designed the experiment, T.P.H collected the data, M.H.R. and T.P.H. analysed the data, M.H.R., and T.P.H. wrote code for data analysis, M.H.R. and T.P.H. wrote the manuscript, M.H.R. and T.P.H. edited the manuscript.

## 5. Supplementary Figures

### 5.1 Spectral power

**Supplementary Figure 1.**
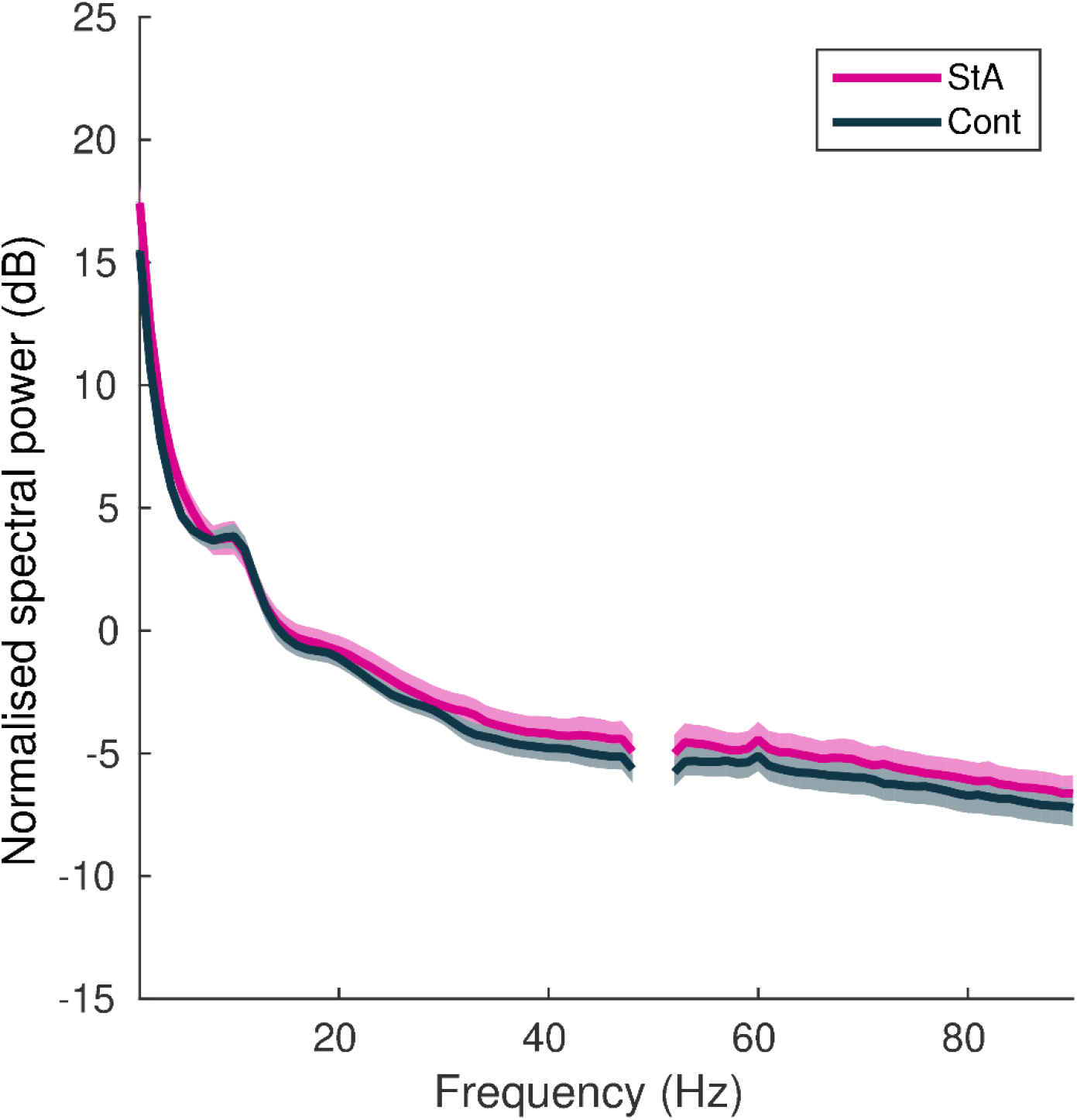
Grand-average of the raw spectral power during task performance. The raw power spectral density during task performance was converted into decibels (dB: 10*log_10_), and grand-averaged separately in state anxious (pink) and control (black) participants. Shaded areas denote the standard error of the mean (SEM). There was no significant difference in raw power between groups (P > 0.05, cluster-based permutation test).

### 5.2 Gamma: pwPE

**Supplementary Figure 2.**
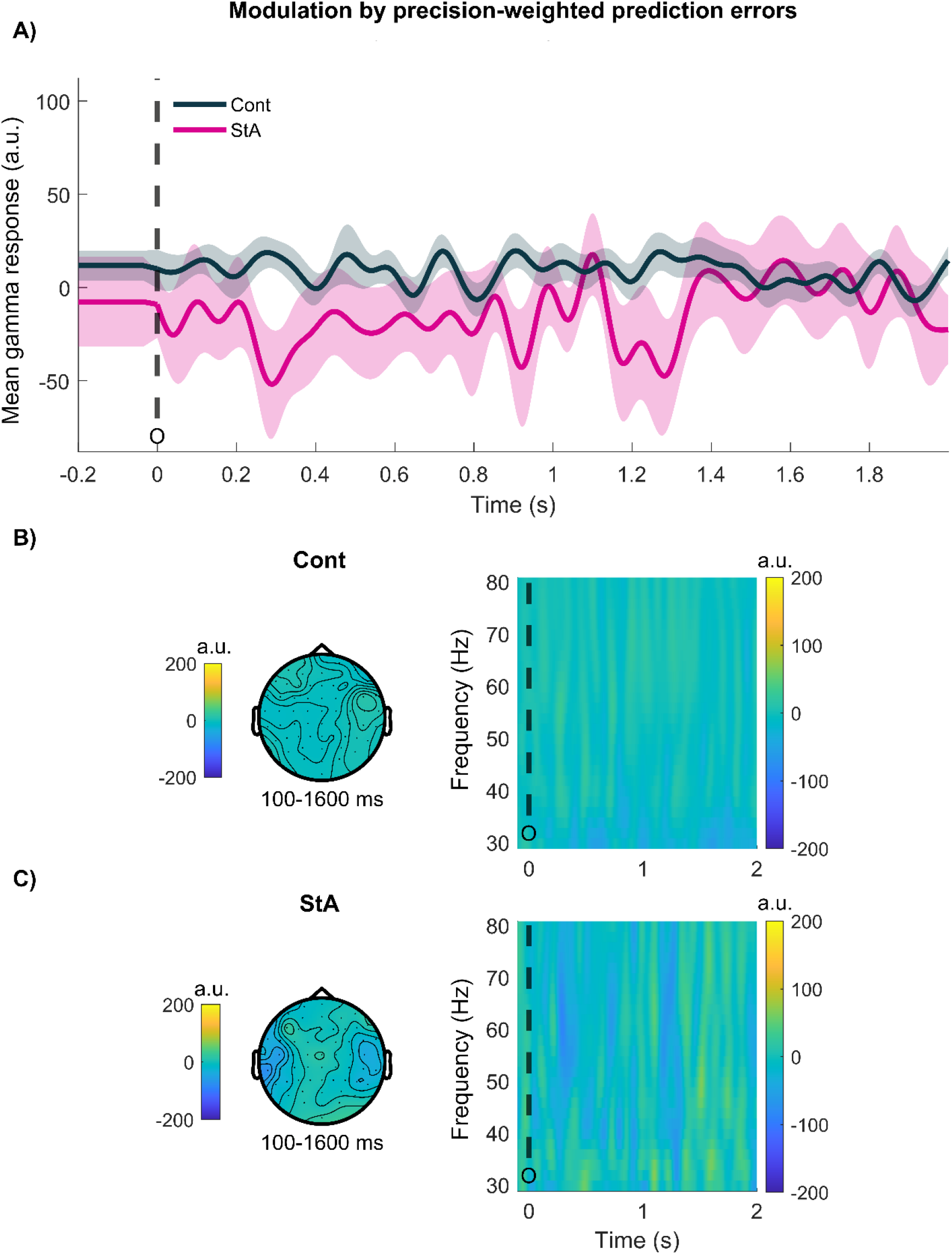
Gamma activity modulated by precision-weighted prediction errors updating belief estimates about the reward tendency. **A)** The average gamma response (31–80 Hz) in arbitrary units (a.u.) to pwPEs on level 2 (|ε_2_|) in each group (Controls, black; StA, pink), with time in seconds (s) on the x-axis. **B)** The correlates of pwPEs |ε_2_| in gamma activity in the Cont group. The left topographic distribution shows activity between 100–1600 ms; the right time frequency image is for |ε_2_| in gamma activity in all electrodes presented 0–2 s from the outcome (black dashed line, ‘O’). **C)** StA topographic representation of |ε_2_| in gamma activity (left), with the time frequency image (right) between 0–2 s (outcome given by black dashed line, ‘O’).

### 5.3 Response regressor: stimulus-locked

**Supplementary Figure 3.**
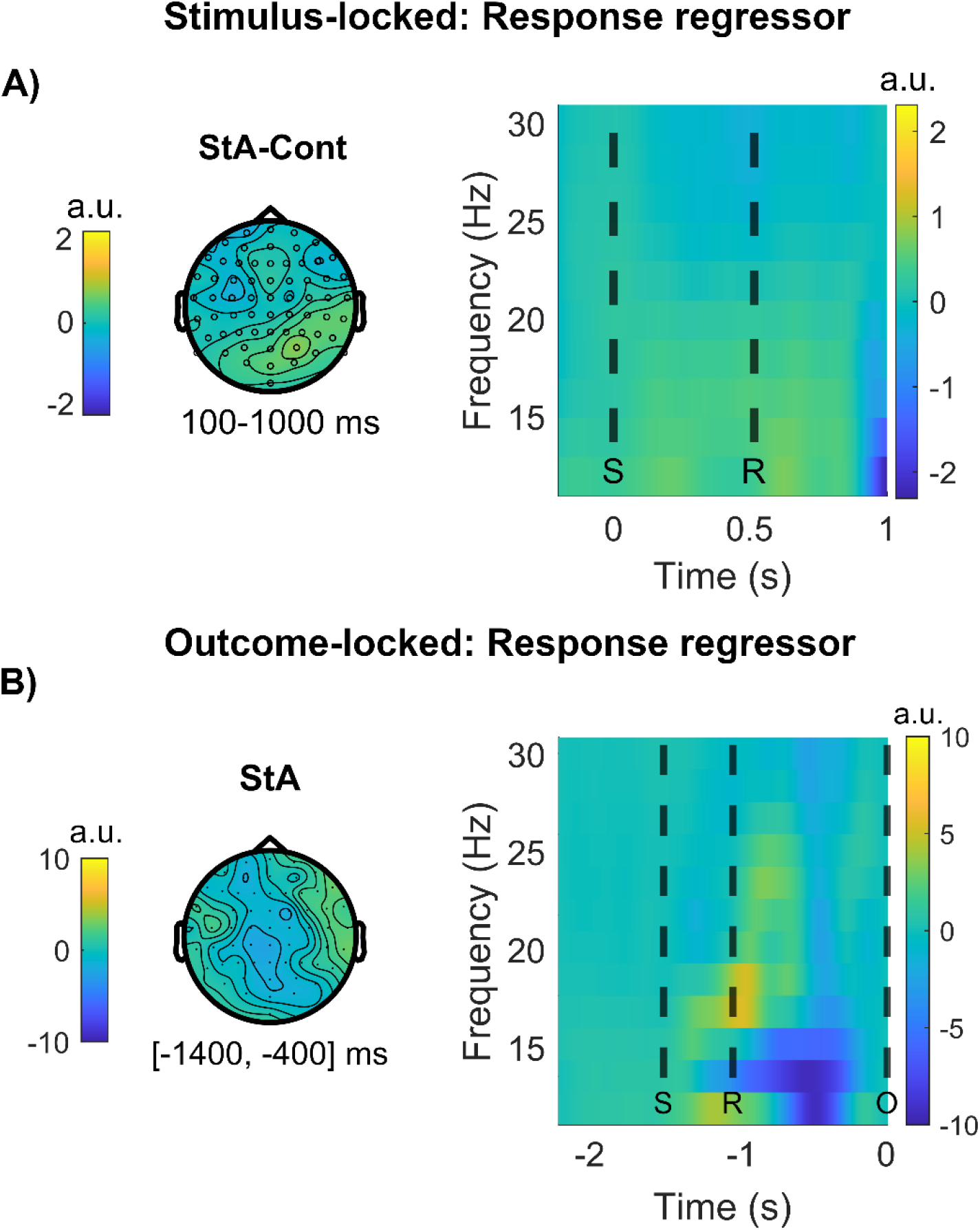
Stimulus and outcome-locked beta (13–30 Hz) changes to response regressor (related to Figure 6). All responses were provided using the right hand. **A)** An independent-samples statistical analysis on stimulus-locked beta activity by the response regressor—in arbitrary units (a.u.)—using cluster-based permutations showed no significant differences between StA and Cont between 100–1000 ms post-stimulus presentation. The right column shows the time-frequency image response in beta activity for the between-group data averaged over all electrodes (0–1 s, stimulus-locked). Dashed black lines represent the average time of the stimuli presentation ‘S’ and the response ‘R’. **B)** For the state anxiety group (StA), we tested the within-group effect of the response regressor on the oscillatory activity locked to the outcome (between −1000 to −100 ms). This analysis showed no significant change in beta activity from baseline. The corresponding time-frequency image to the right shows beta activity averaged across all electrodes (−2 to 0 s, outcome-locked, dashed black lines for the average time of the stimuli presentation ‘S’ and the response ‘R’.

